# TaxoNERD: deep neural models for the recognition of taxonomic entities in the ecological and evolutionary literature

**DOI:** 10.1101/2021.06.08.444426

**Authors:** Nicolas Le Guillarme, Wilfried Thuiller

## Abstract

1. Given the biodiversity crisis, we more than ever need to access information on multiple taxa (e.g. distribution, traits, diet) in the scientific literature to understand, map and predict all-inclusive biodiversity. Tools are needed to automatically extract useful information from the ever-growing corpus of ecological texts and feed this information to open data repositories. A prerequisite is the ability to recognise mentions of taxa in text, a special case of named entity recognition (NER). In recent years, deep learning-based NER systems have become ubiquitous, yielding state-of-the-art results in the general and biomedical domains. However, no such tool is available to ecologists wishing to extract information from the biodiversity literature.
2. We propose a new tool called TaxoNERD that provides two deep neural network (DNN) models to recognise taxon mentions in ecological documents. To achieve high performance, DNN-based NER models usually need to be trained on a large corpus of manually annotated text. Creating such a gold standard corpus (GSC) is a laborious and costly process, with the result that GSCs in the ecological domain tend to be too small to learn an accurate DNN model from scratch. To address this issue, we leverage existing DNN models pretrained on large biomedical corpora using transfer learning. The performance of our models is evaluated on four GSCs and compared to the most popular taxonomic NER tools.
3. Our experiments suggest that existing taxonomic NER tools are not suited to the extraction of ecological information from text as they performed poorly on ecologically-oriented corpora, either because they do not take account of the variability of taxon naming practices, or because they do not generalise well to the ecological domain. Conversely, a domain-specific DNN-based tool like TaxoNERD outperformed the other approaches on an ecological information extraction task.
4. Efforts are needed in order to raise ecological information extraction to the same level of performance as its biomedical counterpart. One promising direction is to leverage the huge corpus of unlabelled ecological texts to learn a language representation model that could benefit downstream tasks. These efforts could be highly beneficial to ecologists on the long term.

## 1 INTRODUCTION

Ecology is rapidly evolving into a data-intensive science that increasingly relies on massive datasets and global knowledge bases to address questions at broader spatial and temporal scales (Michener and Jones, 2012; Soranno and Schimel, 2014; Hallgren et al., 2016; Farley et al., 2018). Although large efforts are being made to elevate research data to be first-class scientific outputs and to promote findable, accessible, interoperable and reusable (FAIR) data (Wilkinson et al., 2016), the overall scientific literature is still a major container for much of the available information on organisms, populations, communities and ecosystems. In addition to the hundreds of millions of pages that make up the historical biodiversity literature, thousands of ecology papers are published every year (Cornford et al., 2020). This represents an enormous amount of unstructured information which is hardly exploitable for large-scale ecological studies, unless we have the tools to automatically extract the relevant information to be fed into open biodiversity databases in a standardized form (Thessen et al., 2012).

Information extraction (IE) is the task of automatically extracting structured information from machine-readable documents, usually from textual corpora expressed in natural language form. Information extraction and its subtasks are key components of a variety of high-level Natural Language Processing (NLP) applications, including knowledge base population (KBP) that is of particular interest for ecologists. KBP consists of discovering new facts about entities from a large corpus of text in order to fill an incomplete knowledge base. In the ecological domain, this includes facts about organism occurrences, phenotypes, habitats, interactions, etc. Such statements are commonly represented in the form of triples (subject, predicate, object), where the subject and the object are entities that have some relationship between them as indicated by the predicate. In most cases, the subject is a taxon, while the object may be a geographical location, the value of a trait measurement, a type of habitat or another taxon depending on the nature of the extracted piece of information. Extracting triples from text to populate a knowledge base is challenging, as it requires the ability to detect mentions of entities of interest in text (= named entity recognition), disambiguate and normalise each textual mention by matching it to the corresponding entity in the knowledge base (= named entity normalisation or disambiguation), and find the semantic relationships that hold between pairs of entities (= relation extraction). A typical information extraction pipeline for knowledge base population is depicted in Fig. 1.

**FIGURE 1.**
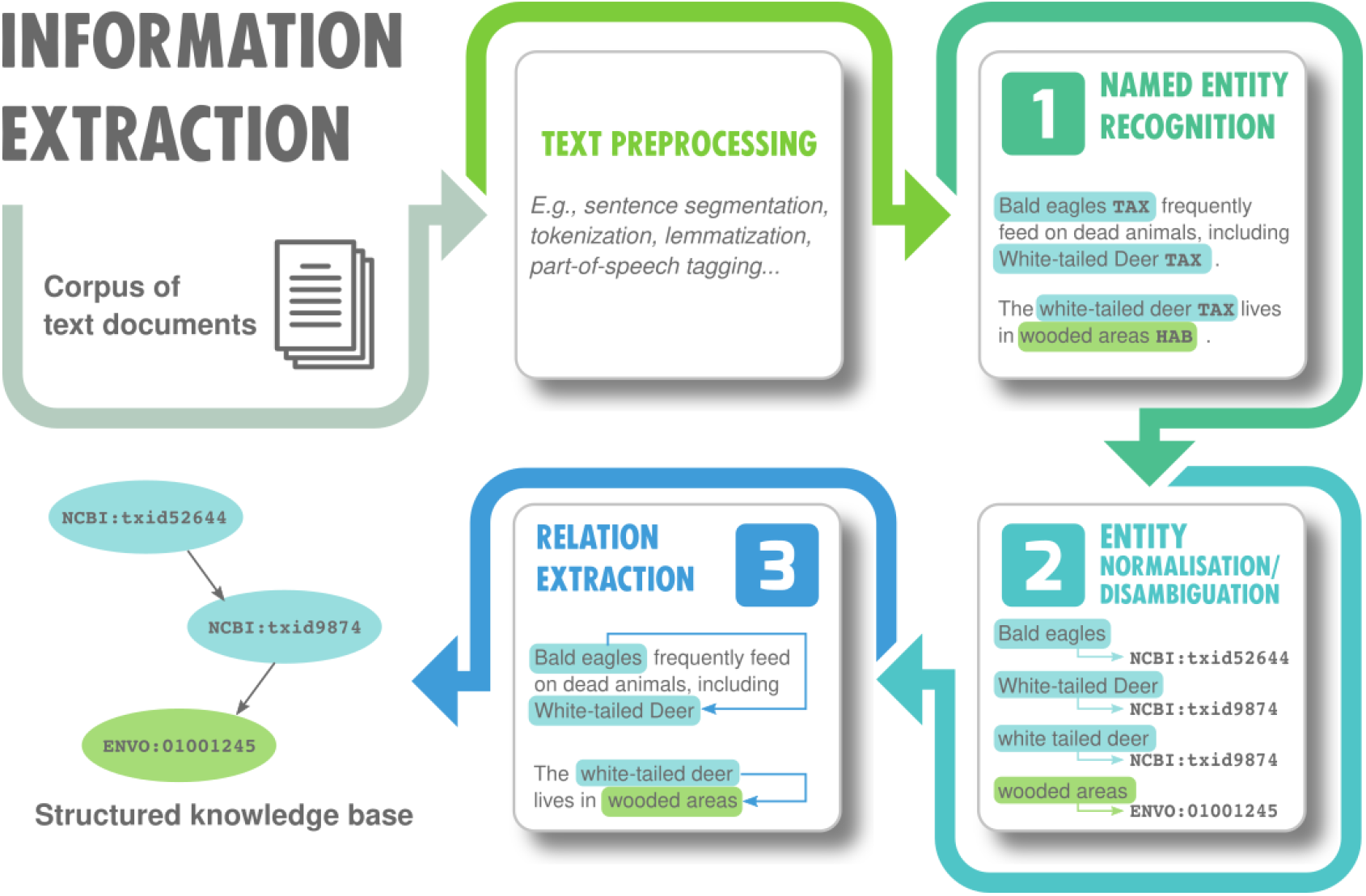
A simple information extraction pipeline for knowledge base population. This pipeline takes a corpus of textual documents as input and generates a collection of factual knowledge represented by triples (entity, relation, entity) as output. This network of interconnected entities forms what is commonly known as a knowledge graph (Ji et al., 2020). Knowledge graphs are a cornerstone of modern artificial intelligence applications.

This paper focuses on the first subtask of information extraction, called *named entity recognition* (NER), and more specifically, on a special case of NER that consists in detecting mentions of taxa in textual documents. Taxonomic entity recognition is critical for augmenting ecological knowledge bases with new facts, as much ecological knowledge refers to some taxonomic unit, whether at the species or at a higher taxonomic level. Identifying taxon names on textual documents is a challenging task. Taxonomic NER systems have to cope with the diversity of taxon naming practices (accepted scientific names with or without authorship information, synonyms, vernacular names in different languages, acronyms and other abbreviations…), the homonymy of some taxon names with common words, and the ambiguity arising from the use of the same common name to refer to distinct species (Gerner et al., 2010).

Previous efforts in taxonomic NER have mainly been directed towards the identification of organism mentions in the biomedical literature. Recognising species and linking them to relevant genes or proteins is indeed critical to the success of many downstream tasks such as gene normalisation and protein-protein interaction extraction (Pafilis et al., 2013). As a consequence, most existing taxonomic NER systems have been designed for biomedical use cases (Gerner et al., 2010; Naderi et al., 2011; Wei et al., 2012; Pafilis et al., 2013; Giorgi and Bader, 2018; Lee et al., 2020), although seminal works focused on the extraction of taxonomic names from biodiversity legacy literature (Koning et al., 2005; Sautter et al., 2006). Several taxonomic NER systems have been developed over the years, using different approaches that can generally be categorised as being based on rules, dictionaries, or machine learning (ML). In addition, a number of tools fall into the category of hybrid systems, combining machine learning with either dictionaries or sets of rules (Naderi et al., 2011; Akella et al., 2012).

Rule-based systems (Koning et al., 2005; Sautter et al., 2006) use handcrafted rules to detect mentions of taxa in text, taking advantage of regularities in taxon naming conventions, e.g. the structure of binomial (Linnean) nomenclature for species names. Consequently, these approaches are more appropriate for detecting scientific names and do not require any updates as taxonomies are revised or new species are discovered. However, they are often unable to identify alternative forms of taxon names such as vernacular names, which do not follow binomial naming conventions, resulting in a low recall. In addition, these methods generally have a low precision as they tend to mistake non-taxonomic scientific terms for taxon names.

Dictionary-based systems (Gerner et al., 2010; Pafilis et al., 2013), on the other hand, are able to recognise taxon names with a high precision by using a well-curated and comprehensive list of taxon names against which chunks of text are matched to identify taxonomic entities. An advantage of dictionary-based approaches over rule-based ones is that they are equally well suited for recognising all types of taxon names. On top of that, entity normalisation is straightforward since dictionaries are generally derived from taxonomic databases such as the NCBI Taxonomy (Federhen, 2002). Although these databases contain a huge number of taxonomic names, they cannot be considered exhaustive as new taxa are continuously described. Therefore, these systems are often characterised by a low recall, as they cannot handle new or abandoned taxon names, misspellings or other unexpected naming variants. In addition, dictionary-based approaches cannot resolve the ambiguity due to homonymy between taxon names and common words as matching is context-agnostic.

Machine learning-based systems replace human-curated rules or fixed lists of names by a statistical model that has been trained to recognise taxon mentions from a feature vector representation of input data (Campos et al., 2012). ML-based systems can be trained to recognise any type of taxon names, depending on whether or not these names have been annotated in the training corpus. ML-based tools are also more robust to new names and misspellings than rule-based and dictionary-based systems. Besides, contextual features can be used to deal with ambiguous names, e.g. homonyms. The main drawback of these approaches is their dependency on annotated documents, which are difficult and expensive to obtain. Furthermore, earlier feature-based ML algorithms rely heavily on handcrafted domain-specific features, requiring considerable engineering skills and domain expertise and leading to highly specialised solutions.

In recent years, deep learning-based NER models have become ubiquitous and have achieved state-of-the-art results in a large number of domains (Li et al., 2020). In particular, remarkable progress has been made in biomedical information extraction through the widespread application of deep learning techniques (Liu et al., 2016; Perera et al., 2020). Deep learning refers to a class of machine learning techniques that use multiple processing layers (typically artificial neural networks) to learn latent representations of data with multiple levels of abstraction (Goodfellow et al., 2016). The ability of deep neural networks (DNNs) to auto-detect hidden features in complex, highly dimensional data removes the burden of task-specific, knowledge-centred feature engineering. In return, their performance largely depends on the availability of large amounts of high quality, manually annotated data in the form of gold standard corpora (GSCs). Indeed, DNNs usually have a large number of parameters, which make them overfit on small training datasets, with the consequence that the resulting models perform poorly on unseen data. However, creating a GSC is laborious and time-consuming, requiring expertise for annotating domain-specific data. As a consequence, GSCs in the ecological domain are few in number and small in size. To tackle the problem of training data shortage, several techniques have been proposed, including data augmentation (Dai and Adel, 2020) and transfer learning (Giorgi and Bader, 2018; Qiu et al., 2020).

Over the last few years, a number of open-source NLP toolkits featuring DNN-based NER solutions have been developed, with an emphasis on accessibility for non-expert users (Dernoncourt et al., 2017; Neumann et al., 2019; Wolf et al., 2019; Giorgi and Bader, 2020). While these toolkits often provide deep models for biomedical NLP, there is so far no such models for ecological applications. As new use cases emerge, including the need to extract factual statements from the ecological literature to augment biodiversity databases with up-to-date information, ecological information extraction sees a resurgence of interest from the community (Tamaddoni-Nezhad et al., 2013; Thessen and Parr, 2014; Compson et al., 2018; Nguyen et al., 2019; Chaix et al., 2019; Muñoz et al., 2019). However, ecologists still lack the tools to build biodiversity information extraction pipelines with state-of-the-art performance.

This paper addresses the task of taxonomic NER as a critical component of such pipelines, and a first step towards the development of a toolkit of state-of-the-art algorithms that would help the ecology and evolution community make the most of the ever-growing corpus of texts on biodiversity. More specifically, we propose a new tool called TaxoNERD^1^ (Taxonomic Named Entity Recognition using DeepModels) that uses deep neural networks to recognise taxonomic entities in the ecological literature. TaxoNERD addresses two challenges of taxonomic NER in these domains: the diversity of taxon naming practices and related problems (e.g. homonymy, ambiguity or variability), and the relatively small size of the few GSCs available for this task. It is our hope that making such tools accessible for ecologists and practitioners will pave the way towards the development of new tools for ecological information extraction and their wider adoption by the community.

## 2 MATERIALS AND METHODS

The following sections introduce the two network architectures used to train TaxoNERD’s taxonomic NER models, as well as the pretrain-and-finetune approach adopted to learn these models from an ecological gold standard corpus. We also describe the corpora and metrics used for evaluation, and briefly present the existing NER tools against which we compared our approach.

### 2.1 TaxoNERD’s model architectures

#### 2.1.1 spaCy’s NER model

spaCy^2^ is an increasingly popular open-source library for advanced Natural Language Processing in Python. spaCy provides a variety of practical tools to build information extraction or natural language understanding systems, including pretrained DNN models for named entity recognition, part-of-speech tagging, dependency parsing, text classification and more. spaCy’s models have emerged as the *de facto* standard for practical NLP due to their speed, robustness and close to state of the art performance. In addition, spaCy makes it easy to create, train, manage, deploy and use custom NLP pipelines. For all these reasons, we choose to build upon the spaCy library to create our taxonomic NER system.

spaCy’s NER models rely on a pretty generic neural architecture, depicted in Fig. 2. This architecture consists of two subnetworks. The first subnetwork learns an embedding model whose role is to embed tokens (≈words) into a continuous vector space. A word embedding is a low-dimensional real-valued vector representation of a word (Zhang et al., 2016). Word embeddings encode the meaning of the words they represent in the sense that the words that are closer in the vector space are expected to be similar in meaning. There are two kinds of word embeddings (Qiu et al., 2020): non-contextual and contextual embeddings.

**Non-contextual embeddings.** A word is mapped to a single context-independent vector representation using a lookup table. This lookup table is usually learned from a large corpus of unlabelled text using self-supervision (Mikolov et al., 2013; Pennington et al., 2014). One of the main limitations of non-contextual word embeddings is that words with multiple meanings are conflated into a single representation. Therefore, these embeddings cannot handle polysemy (one word having multiple meanings) and homonymy (words that share the same spelling but with different meanings) properly. Another issue is the out-of-vocabulary problem: models can only produce meaningful embeddings for words that have been seen in the training corpus.
**Contextual embeddings.** To address the issue of polysemy and the context-dependent nature of words, contextual word embeddings move beyond word-level semantics in that each token is associated with a representation that is a function of the entire input sequence, thereby capturing uses of words across varied context. Contextual embeddings are typically obtained by mapping each input token in the sequence to its non-contextualised representation first, before applying an aggregation function to encode context. This aggregation function is usually modelled by a deep neural network, which is then called a neural contextual encoder. There are many possible architectures for this encoder (see Li et al. (2020) and Qiu et al. (2020) for a survey). Contextual embeddings pre-trained on large-scale unlabelled corpora achieves state-of-the-art performance on a wide range of NLP tasks.

**FIGURE 2.**
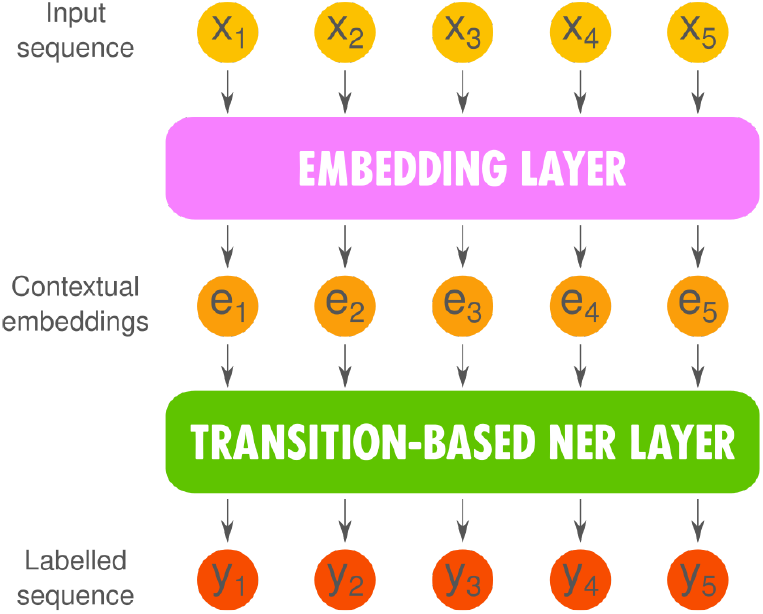
spaCy’s generic neural architecture for named entity recognition.

The second subnetwork assigns class labels to non-overlapping spans of tokens using a probabilistic transition-based chunking model similar to Lample et al. (2016). This model relies on a stack data structure to incrementally construct chunks of the input sequence and assign a class label to those chunks that correspond to named entities. At each time step, the possible actions (add a token to the stack, assign a label to the current chunk…) are scored by feeding a representation of the current state of the stack to a multilayer neural network. This representation is obtained by combining the embeddings of the tokens that make up the stack. Then the action with the highest score is chosen and the stack moves to another state. The process is repeated until the algorithm reaches a termination state.

#### 2.1.2 TaxoNERD’s NER models

TaxoNERD offers the user a choice of two NER models, with a different balance between speed and accuracy. The two models use the same two-layer architecture as spaCy’s NER models but differ in the architecture of their embedding layer.

##### en_ner_eco_md: a model designed for speed

The en_ner_eco_md model (Fig. 3a) is a taxonomic NER model that uses spaCy’s standard Tok2Vec layer to generate contextual embeddings for the input tokens. This embedding layer is itself composed of two subnetworks.

**FIGURE 3.**
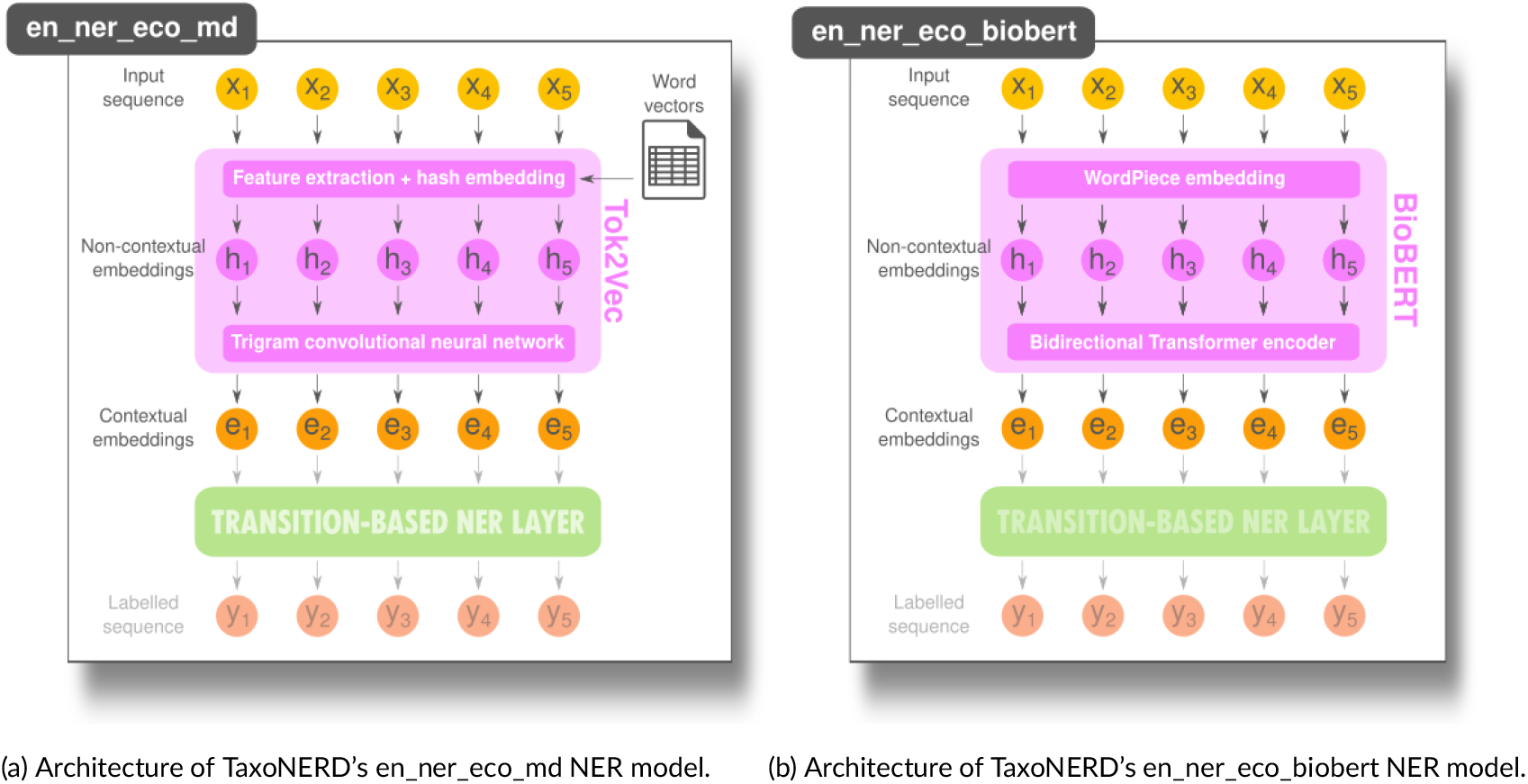
TaxoNERD’s deep NER models differ in the architecture of their embedding layer. The en_ner_eco_model combines hash embeddings and CNN-based contextual encoding for speed, while the en_ner_eco_biobert model leverages a Transformer-based pretrained language model called BioBERT for accuracy.

The first subnetwork embeds input tokens into context-independent word vectors. The model first extracts a number of subword features (normalised string form, prefix, suffix and word shape), each of which is embedded separately using hash embedding (Svenstrup et al., 2017). Subword representations are concatenated and the resulting vector is passed through a feed-forward subnetwork to generate a word vector for the input token. Enriching word vectors with subword information is a common approach for tackling the out-of-vocabulary problem (Bojanowski et al., 2017). In addition to subword features, spaCy’s standard embedding layer can use pretrained word vector tables as additional features, which sometimes results in significant improvements in accuracy. The en_ner_eco_md model uses a 50k word vector table trained on a biomedical corpus and provided as part of scispaCy^3^, a Python library for practical biomedical/scientific text processing which heavily leverages the spaCy library (Neumann et al., 2019).

The second subnetwork encodes context into the context-independent embeddings generated by the first subnetwork using a convolutional neural network (CNN). CNNs are a class of deep neural networks, most commonly applied to image processing and computer vision, that uses a series of convolution layers to aggregate local information from multiple pixels/words/… and generate low-dimensional representations of the input data that successfully captures the spatial and/or temporal dependencies (Brodrick et al., 2019). The basic building block of spaCy’s contextual encoder consists of a 3-gram convolution layer, that basically concatenates the vector representations of a token and its two neighbours, followed by a multi-layer perceptron that maps this concatenated vector to a lower dimensional output vector. This whole block (3-gram convolution layer + multi-layer perceptron) enables to relearn the meaning of a word (i.e. its embedding) based on its direct neighbours. By stacking more such blocks in the CNN architecture, the size of the surrounding context used to recalculate the embedding of a word increases, thus incorporating more contextual information in the representation.

The NER model resulting from the combination of CNN-based word embedding and transition-based sequence labelling is an efficient alternative to the standard solutions based on recurrent neural networks (RNNs) which have long dominated the NLP landscape (Lample et al., 2016; Giorgi and Bader, 2018; Li et al., 2020) and are now gradually being deposed by the Transformer model (Vaswani et al., 2017). In particular, spaCy’s NER model is smaller and computationally cheaper. It therefore runs much faster than these state-of-the-art deep models, while delivering close performance.

##### en_ner_eco_biobert: a model designed for accuracy

Since version 3.0, spaCy has added support for Transformer models. Transformers (Vaswani et al., 2017) are a family of neural network architectures that utilise the mechanism of self-attention, i.e. weighing the influence of different parts of the input sequence, to capture long-range dependencies in sequential data. Transformers allow for significantly more parallelisation than sequence models like CNNs and RNNs, and therefore reduced training times. Due to this feature, Transformers have rapidly become the mainstream architecture for many NLP problems, replacing older RNN models such as the long short-term memory (LSTM), and bringing NLP to a new era.

Transformers are now commonly used to pretrain language representation models from a large amount of unannotated text. In contrast to GSCs, large-scale unlabelled corpora are relatively easy to construct. Such corpora can be leveraged by learning contextualized word representation models from them in an unsupervised manner. Then, with minimal architectural modification, the resulting pretrained language model can be applied to various downstream NLP tasks via a procedure called transfer learning Giorgi and Bader (2020). The use of word embeddings extracted from pretrained language models has brought significant performance gains on a number of NLP tasks, including named entity recognition. Hundreds of pretrained language models based on the Transformer architecture are now available through libraries such as HuggingFace’s Transformers (Wolf et al., 2019), including general purpose language representation models such as BERT (Devlin et al., 2018) and XLNet (Yang et al., 2019), and domain-specific language models, such as SciBERT (Beltagy et al., 2019) for scientific text processing and BioBERT (Lee et al., 2020) for biomedical text mining. To our knowledge, there is no pretrained language model for ecological or evolutionary applications.

In the en_ner_eco_biobert NER model (Fig. 3b), spaCy’s standard Tok2Vec embedding layer is replaced by BioBERT (Bidirectional Encoder Representations from Transformers for Biomedical TextMining), a domain-specific language representation model pretrained on large-scale biomedical corpora (PubMed abstracts and PMC full-text articles). In the absence of a Transformer model pretrained on ecological corpora, we chose a language model whose domain has commonalities with ecology and evolution, and which shares with them a number of common entities of interest, notably taxa. In addition, Lee et al. (2020) obtained state-of-the-art results in a number of biomedical NLP tasks, including named entity recognition, relation extraction, and question answering, using BioBERT word vector representations. A recent survey also showed that of 6 open-source language models, BioBERT performed best on biomedical tasks Lewis et al. (2020).

Large pretrained Transformer models are tremendously effective for many NLP tasks. However, they have two main limitations. First, they usually require a large training corpus and easily overfit on small or modestly-sized datasets. Although large-scale unlabelled corpora are far easier to obtain than large-scale GSCs, building a corpus large enough (several billion words) to learn such a model requires significant engineering efforts. Second, inference in large Transformer models is prohibitively slow and expensive due to the use of self-attention in multiple layers. Therefore, the en_ner_eco_biobert model is more suitable for use cases where accuracy is more important than computation time.

### 2.2 Training TaxoNERD’s models using transfer learning

Performance of DNN-based approaches to NER largely depends on the availability of large amounts of high quality, manually annotated data in the form of gold standard corpora. The deeper the network architecture, the better the expected performance, but a much larger dataset is needed to fully train model parameters and prevent overfitting. In domains where GSCs tend to be small, as is the case in ecology and evolution, training such large neural networks from scratch, starting with randomly initialised weights, would overfit the training set badly, which would cause the resulting models to perform poorly on unseen data.

One approach to get around this problem is to first pretrain a DNN on a source task for which a large dataset is available. Then, the pretrained weights of this network are used to initialise the weights of a second network, which we continue to train on our typically smaller dataset for the target task (Fig. 4). This process, called transfer learning, has been shown to improve generalisation of the model, reduce training time on the target dataset, and reduce the amount of labelled data needed to obtain high performance (Giorgi and Bader, 2018). There are basically two common ways to transfer knowledge learned from one task or domain to another: feature extraction and fine-tuning.

**Feature extraction.** The pretrained model is used as an off-the-shelf feature extractor. The pretrained weights of the feature extraction layer are frozen, and a new classification layer is trained on the target dataset.
**Fine-tuning.** All the pretrained network’s weights are unfrozen and updated (fine-tuned) for the target task.

**FIGURE 4.**
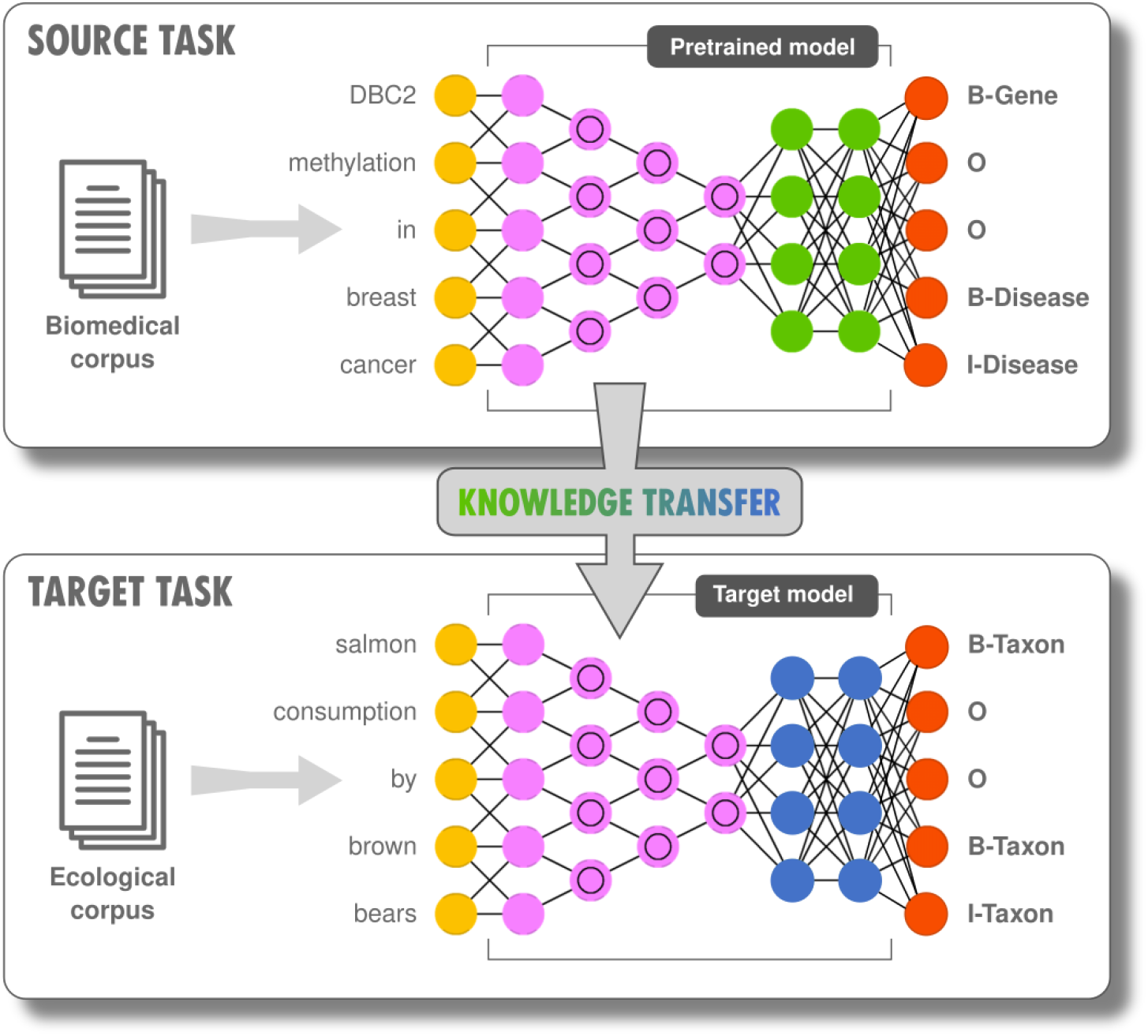
State-of-the-art NER systems are based on deep neural networks that learn latent features from large amounts of data. When only small datasets are available for the target task, a common approach is to use transfer learning. In this example, transfer learning is used to adapt a DNN trained on a large biomedical corpus to the ecological domain. The feature extraction subnetwork (pink nodes) is frozen, while the NER layers (green, then blue nodes) are retrained on the ecological corpus. Alternatively, the pretrained model parameters can be unfrozen and the whole network be fine-tuned on the target task corpus.

The choice between feature extraction and fine-tuning may be guided by some criteria such as the size of the target dataset, and the similarity between the source and target datasets. In practice, fine-tuning is usually more general and convenient for many different downstream tasks, while requiring minimal architecture modifications.

Following the pretrain-and-finetune approach that has become the dominant paradigm for NLP applications in the last few years, we trained TaxoNERD models by reusing embedding layers that have been pretrained on large biomedical corpora, and fine-tuning the pretrained models on a GSC which was built by combining the COPIOUS and Bacteria Biotope task corpora. More precisely, the en_ner_eco_md model is trained by fine-tuning the en_sci_core_md model provided by the scispaCy library. As this model already includes a transition-based NER layer (trained to recognise biomedical entities in general, without distinction between different types of entities), we kept the NER layer as it is, simply added a new class of entities corresponding to taxon names, and fine-tuned the whole network (embedding and labelling subnetworks) on our ecological GSC. To learn the en_ner_eco_biobert model, we used the BioBERT language representation model as our embedding layer, followed by a transition-based NER layer with one output class (taxon name) and randomly initialised weights, and fine-tuned the whole network on our GSC. In both cases, all the layers are updated for the target task.

The networks for the target task were fine-tuned using Adam optimisation, with standard parameter values. The batch size was increased from 1 to 32 during training, as it has shown to be an effective trick (Smith et al., 2017). For regularisation, dropout was set to 0.2, and early stopping was used on the validation set with a patience of 1000 steps, i.e. the model stopped training if performance did not improve on the validation set during the last 1000 iterations (contrary to most deep learning framework, patience in spaCy is not specified as a number of epochs but as a number of steps).

### 2.3 TaxoNERD’s models evaluation

#### 2.3.1 Gold standard corpora for taxonomic NER

We evaluated TaxoNERD’s models on 4 gold standard corpora specifically designed for taxonomic NER or with a strong focus on taxon names recognition : LINNAEUS, Species-800, COPIOUS, and the Bacteria Biotope task corpus (see Table 1 for a summary of these corpora). All four corpora are in English. Annotations usually include the boundaries of the named entity (start and end character offsets), its class and the entity’s text, and are written in some annotation format, the two most common being the Standoff and IOB2 formats (Fig. 5). To facilitate evaluation, all annotations were converted to the Standoff format.

**FIGURE 5.**
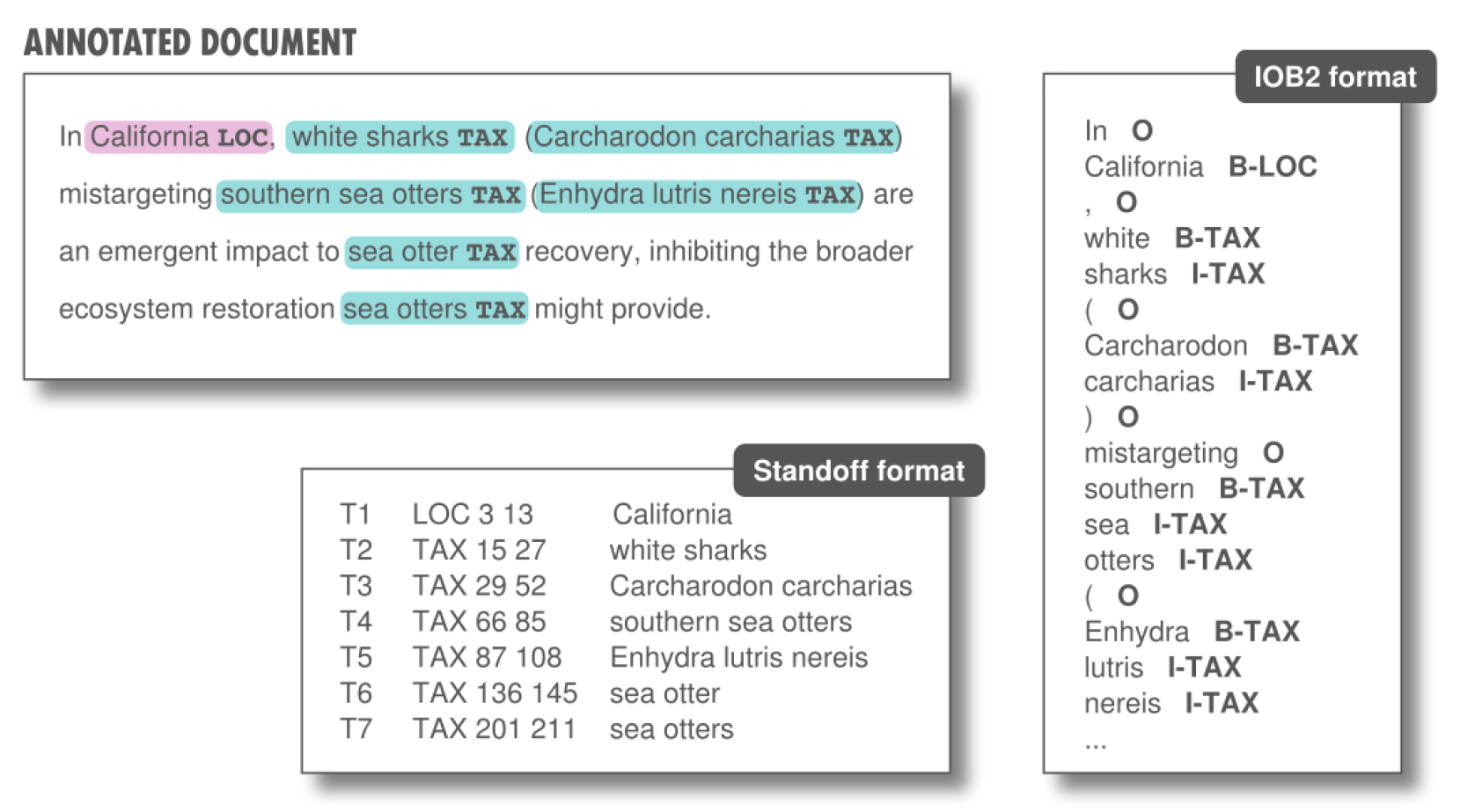
A gold standard corpus is a collection of manually annotated documents. An entity’s annotation includes at least the left and right boundaries of the entity span as well as its class. Standoff and IOB2 are the two most common tagging formats. In IOB2, the B-prefix indicates that the token is the beginning of an entity, and the I-prefix indicates that the token is inside an entity. An O tag indicates that a token belongs to no entity.

**TABLE 1.** Gold standard corpora (GSCs) used for evaluation.

| Corpus | Documents | Taxon mentions | NCBI IDs | Taxonomic rank | Documents type |
| --- | --- | --- | --- | --- | --- |
| LINNAEUS | 100 | 4,259 | Y | Species | PubMed Central full papers |
| S800 | 800 | 3,708 | Y | All | MEDLINE abstracts |
| COPIOUS | 668 | 12,227 | N | All but microorganisms | Biodiversity Heritage Library pages |
| BB task corpus | 392 | 2,487 | Y | Only microorganisms | PubMed abstracts + full-text extracts |

LINNAEUS (Gerner et al., 2010) is a GSC of 100 full-text biomedical articles that were randomly selected from the open-access subset of PubMed Central and manually annotated for species mentions. Mentions of genera or other higher-order taxonomic ranks (family, order, class…) were not annotated since it was not the focus of the original work. After annotation, all mentions of species terms were normalised by matching each mention to the corresponding taxon in the NCBI Taxonomy (Federhen, 2002). Of the 4,259 species mentions annotated in this corpus, 72% are common names, including terms that do not directly convey species names, such as “*patient*”, “*child*”, “*boy*” which indirectly refer to subspecies *Homo sapiens sapiens*. A total of 65% of all mentions are normal species mention while 28% are adjectival modifiers (e.g. “*human*” in “*human P53*”).

Species-800 (Pafilis et al., 2013), or S800, is a GSC that was developed to increase the diversity of species names compared to the LINNAEUS corpus. S800 was constructed by randomly selecting 100 MEDLINE abstracts from journals in each of the following 8 categories: bacteriology, botany, entomology, medicine, mycology, protistology, virology, and zoology. Taxon mentions, including Linnaean binomials, common names, strain names, and author-defined acronyms, were manually annotated. While the main focus was on annotating species mentions, other taxonomic ranks (e.g. kingdoms, orders, genera, strains) were also considered. The S800 corpus contains approximately the same number of annotated species mentions as the LINNAEUS corpus (3,708 mentions). However, the former contains more than three times as many unique species and names as the latter.

COPIOUS (Nguyen et al., 2019) is a GSC directed towards the extraction of species occurrence from the biodiversity literature. As such, COPIOUS covers a wider range of entities than LINNAEUS and S800, including taxon names, geographical locations, habitats, temporal expressions and person names. The COPIOUS corpus consists of 668 document pages downloaded from the Biodiversity Heritage Library (BHL). More than 28K entities have been manually annotated by experts, 44% (12,227) of which are taxa. Annotated taxon mentions include species, genera, families and all higher-order taxonomic ranks. Both current and historical scientific names were annotated. For scientific names that include authorship information, two entities were created, one with the authorship information, the other without. These entities are overlapping as they share a common substring. Annotations also include vernacular names of species but vernacular names of taxonomic classes (e.g. fish, birds, mammals…) were not tagged as taxon names. Since COPIOUS was developed specifically for extracting information about Philippine biodiversity, a non-negligible part of common names are English transcriptions of Filipino names. However, the authors state that the corpus is general enough to be employed for other biodiversity applications. Also, all microorganism names were excluded as COPIOUS focuses on highly endangered species.

Conversely, the Bacteria Biotope (BB) task corpus (Bossy et al., 2019) focuses on the extraction of information about microbial ecology. This GSC comprises 215 PubMed abstracts related to microorganisms and 177 extracts of variable lengths (from one single sentence to a few paragraphs) selected from 20 full-text articles about microorganisms of food interest. All mentions of microorganism names, habitats and phenotypes have been manually annotated, as well as mentions of geographical places. In addition, microorganisms are normalised to taxa from the NCBI Taxonomy, and habitat and phenotype entities are normalised to concepts from the OntoBiotope ontology. Mentions of microorganisms represent 34% (2,487) of the 7,232 entity mentions in the corpus. 54.8% of these microorganism mentions have no exact string match with any concept in the NCBI Taxonomy.

Both COPIOUS and the BB task corpus contain overlapping entities, which are not supported by TaxoNERD’s NER models or any of the other evaluated systems. To get rid of overlapping entities, we preprocessed the entire corpora by replacing all overlapping entities by the entity corresponding to their union, thus favouring longer mentions (for instance, scientific names with authorship over simple binomial names).

#### 2.3.2 NER evaluation metrics

Each GSC was split into three disjoint subsets : one for training, one for validation during training, and one for testing. Only the test set was used during evaluation. Although the training and validation sets of LINNAEUS and S800 were not used for learning TaxoNERD’s models, we decided to evaluate the methods on their test sets only so that results are easier to compare with those obtained by models trained using these corpora (like the species recognition models from Giorgi and Bader (2018) and Lee et al. (2020)). For LINNAEUS, we used the train/validation/test split of Giorgi and Bader (2018). For S800, we used the split_s800 script^4^ to generate the three subsets. COPIOUS train, test and validation subsets are available on the COPIOUS project webpage^5^. Finally, since the BB task corpus has been published as part of a BioNLP challenge^6^, annotations are provided only for the train and validation sets. Therefore, we used the validation set for testing, and randomly split the train set into train/validation subsets with a 85:15 ratio.

Precision, recall and F-score are commonly used to assess and compare NER systems using gold standard corpora. Precision is the percentage of predicted entities that match gold entities (i.e. entities that are annotated in the GSC), recall is the percentage of gold entities that are correctly predicted, and F-score (also called F_1_-score) combines these two measures into a single score and is defined as the harmonic mean of precision and recall. Whether a prediction is considered correct depends on the matching criterion used. The most common criterion is exact match: a predicted entity is counted as a true positive if both its boundaries and its class fully coincide with a gold entity (see Fig. 6).

**FIGURE 6.**
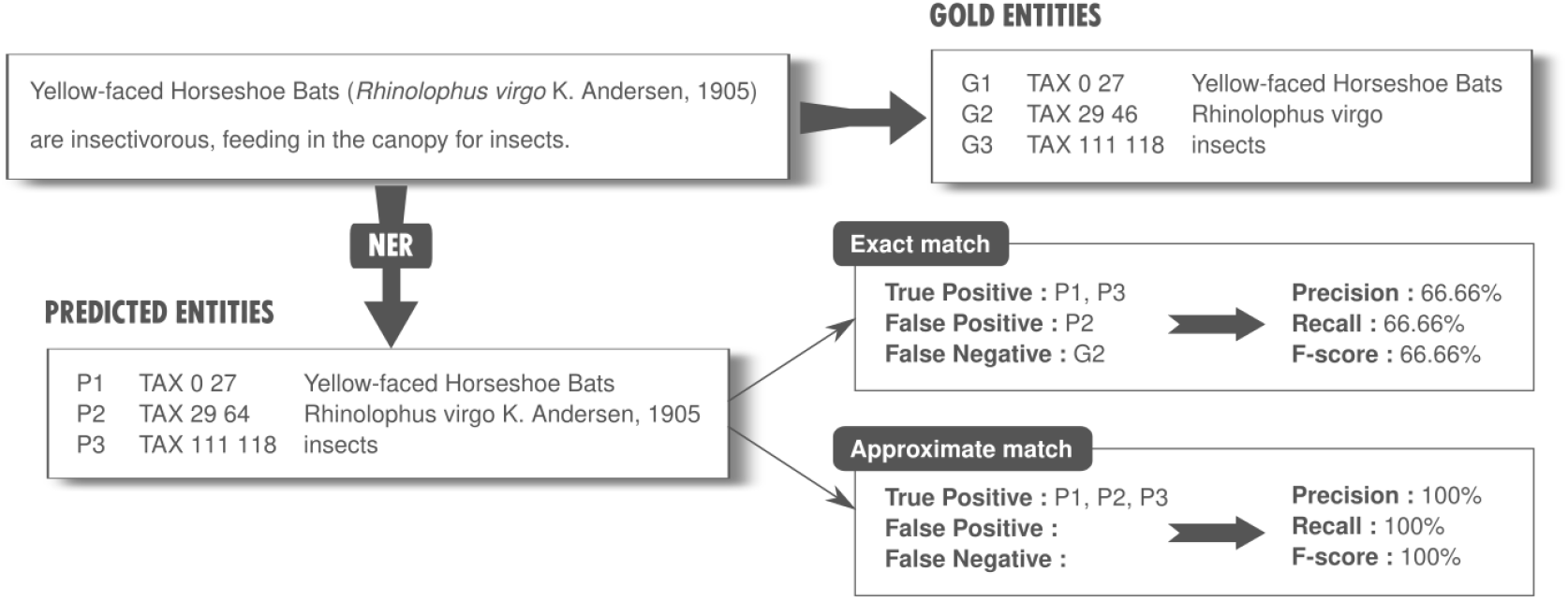
All methods were evaluated in terms of precision, recall and F-score using two matching criteria: exact and approximate match. As the exact match criterion tends to be too strict, underestimating the performance of NER methods in practical settings, we also used a relaxed criterion that judges as correct a predicted entity if it encompasses a gold entity.

However, exact match may not be the most appropriate criterion for evaluating taxonomic NER systems. The annotation of entity boundaries in a GSC depends on the task the corpus was designed for, but also on the person performing the annotation. For example, annotation guidelines may ask the annotator to include authorship information in scientific names or to stick to the binomial name only. Sometimes, both versions are annotated and we end up with two overlapping entities (as in COPIOUS). Using exact match, a NER system that was designed to detect taxon names with authorship will exhibit lower performance on a corpus in which only binomial names were annotated, although it is able to find the relevant piece of information. Annotation inconsistencies are also very frequent within a corpus. Inter-annotator disagreement may be due to different interpretations of annotation guidelines, or to difference in the level of expertise of each annotator. Inconsistencies also exist in the work of a single annotator, as it has often been observed that the annotation behaviour changes over time as annotators gain more experience with their task (Leser and Hakenberg, 2005).

Irregularities in the annotation scheme tend to underestimate NER methods performance, as these methods may correctly predict an entity without exactly matching the gold entity boundaries. This is punished twice by the exact match criterion, since it results in both a false negative and a false positive for simultaneously missing the gold entity and predicting a partial match. One possible solution is to relax the matching criterion to a certain degree Tsai et al. (2006). Indeed, in many applications, finding pieces of information is better than finding nothing at all, and exact match may not reflect the true performance of a system in a practical setting. In addition to exact match, we evaluated all methods using a relaxed criterion, approximate match, that counts a predicted entity as a true positive if there is some gold entity that is a substring of the predicted entity (see Fig. 6). This criterion reflects the fact that for information extraction applications, it is often better to overestimate entity boundaries than to miss relevant information.

### 2.4 TaxoNERD: comparison with existing approaches

We compared TaxoNERD’s NER models en_ner_eco_md and en_ner_eco_biobert with the most popular taxonomic NER systems currently available to ecologists and practitioners. We chose to include only those tools that are readily available either as standalone command-line tools or as high-level libraries that can be easily reused to build complex information extraction pipelines. A summary of the features of each evaluated tool is provided in Table 2.

**TABLE 2.**
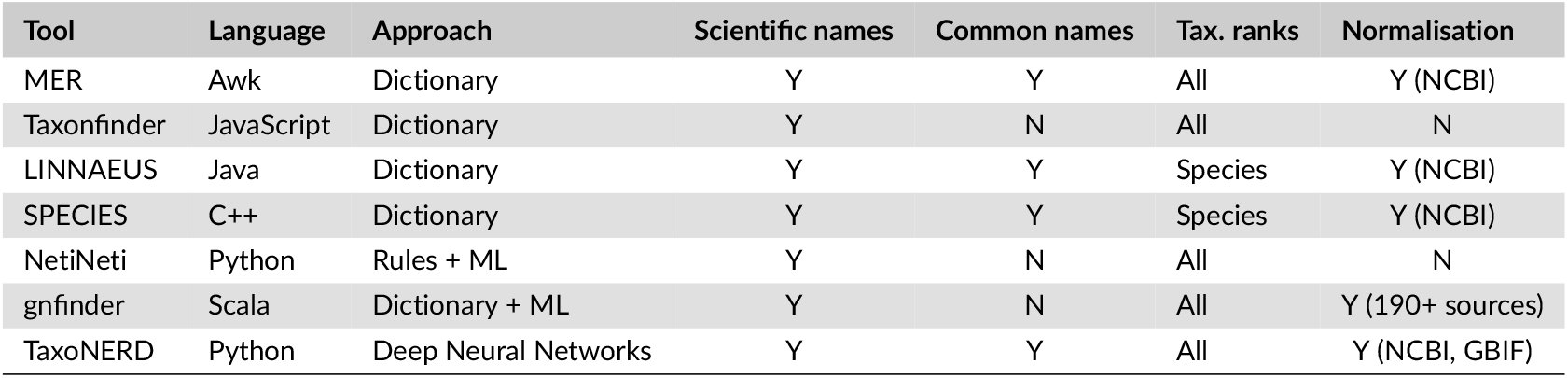
NER tools selected for evaluation.

| Tool | Language | Approach | Scientific names | Common names | Tax. ranks | Normalisation |
| --- | --- | --- | --- | --- | --- | --- |
| MER | Awk | Dictionary | Y | Y | All | Y (NCBI) |
| Taxonfinder | JavaScript | Dictionary | Y | N | All | N |
| LINNAEUS | Java | Dictionary | Y | Y | Species | Y (NCBI) |
| SPECIES | C++ | Dictionary | Y | Y | Species | Y (NCBI) |
| NetiNeti | Python | Rules + ML | Y | N | All | N |
| gnfinder | Scala | Dictionary + ML | Y | N | All | Y (190+ sources) |
| TaxoNERD | Python | Deep Neural Networks | Y | Y | All | Y (NCBI, GBIF) |

LINNAEUS^7^ (Gerner et al., 2010) and SPECIES^8^ (Pafilis et al., 2013) are two popular dictionary-based command-line softwares for taxon names recognition in biomedical documents. LINNAEUS dictionary of names covers 386,108 species and 116,557 higher-order taxonomic ranks, while SPECIES dictionary contains all the species and strain names from the NCBI Taxonomy (as of 2013), including scientific names, common names and other synonyms. Both tools also include abbreviations that were generated automatically from species scientific names (for instance *D. melanogaster* from *Drosophila melanogaster*).

Taxonfinder^9^ (Leary, 2014), NetiNeti^10^ (Akella et al., 2012) and Global Names Finder^11^ (gnfinder) all belong to the category of scientific name taggers as they were specifically designed to recognize mentions of scientific names only. Taxonfinder uses a combination of regular expressions and dictionaries to tag organism scientific names in text. Taxonfinder maintains separate dictionaries for species, genera and higher-order taxonomic ranks, all derived from a manually curated version of NameBank^12^. NetiNeti is a hybrid rule-based/machine learning solution to recognise scientific names of organisms in biomedical and biodiversity literature, including misspelled and new species names. Candidate mentions are identified using simple scientific name capitalisation and abbreviations rules and fed to a binary classifier (Naïve Bayes) to decide whether they are a scientific name or not. Gnfinder is a hybrid dictionary-based and machine learning system for scientific names detection in text. Since December 2019, it has replaced TaxonFinder and NetiNeti as the name-finding engine in the Global Names Recognition and Discovery service (Mozzherin and Shorthouse, 2019). It is currently used by the Biodiversity Heritage Library to locate taxonomic names in their corpus of legacy biodiversity documents (Constantino, 2020). Gnfinder uses a set of dictionaries (including separate dictionaries for species, genera and uninomials) to detect scientific name candidates and extract a number of useful features that are fed to a Naïve Bayes classifier to refine predictions.

We deliberately discarded taxonomic NER tools that cannot be considered as standalone tools because they are tied to a specific NLP architecture, e.g. the OrganismTagger (Naderi et al., 2011) system which comes as a GATE pipeline, or because they are only available as RESTful APIs, e.g. SR4GN (Wei et al., 2012). We also ruled out software that were not or very poorly documented, e.g. TaxonGrab (Koning et al., 2005), although we decided to include Taxonfinder because it is quite self-explained. Because LINNAEUS and SPECIES are tied to their built-in lists of names, specifically designed for biomedical use cases, their results on ecological corpora may not be representative of the true predictive power of dictionary-based approaches. We decided to include, as a baseline, our own dictionary-based taxonomic NER system created using MER (Couto and Lamurias, 2018), a minimal named-entity recognition and linking tool which only requires a lexicon with the list of terms representing the entities of interest. We created this lexicon by extracting all the taxon names from a dump of the NCBI Taxonomy, for a total of about 3.4M names.

## 3 RESULTS

All scripts used for methods evaluation are available on the accompanying GitHub page^13^. TaxoNERD’s models ability to detection mentions of taxa in text was evaluated on all four GSCs and compared to five existing taxonomic NER systems and one dictionary-based baseline. Performance are measured in terms of precision, recall and F-score using both exact and approximate match criteria. The results are presented in Table 3 and shown in Fig. 7.

**FIGURE 7.**
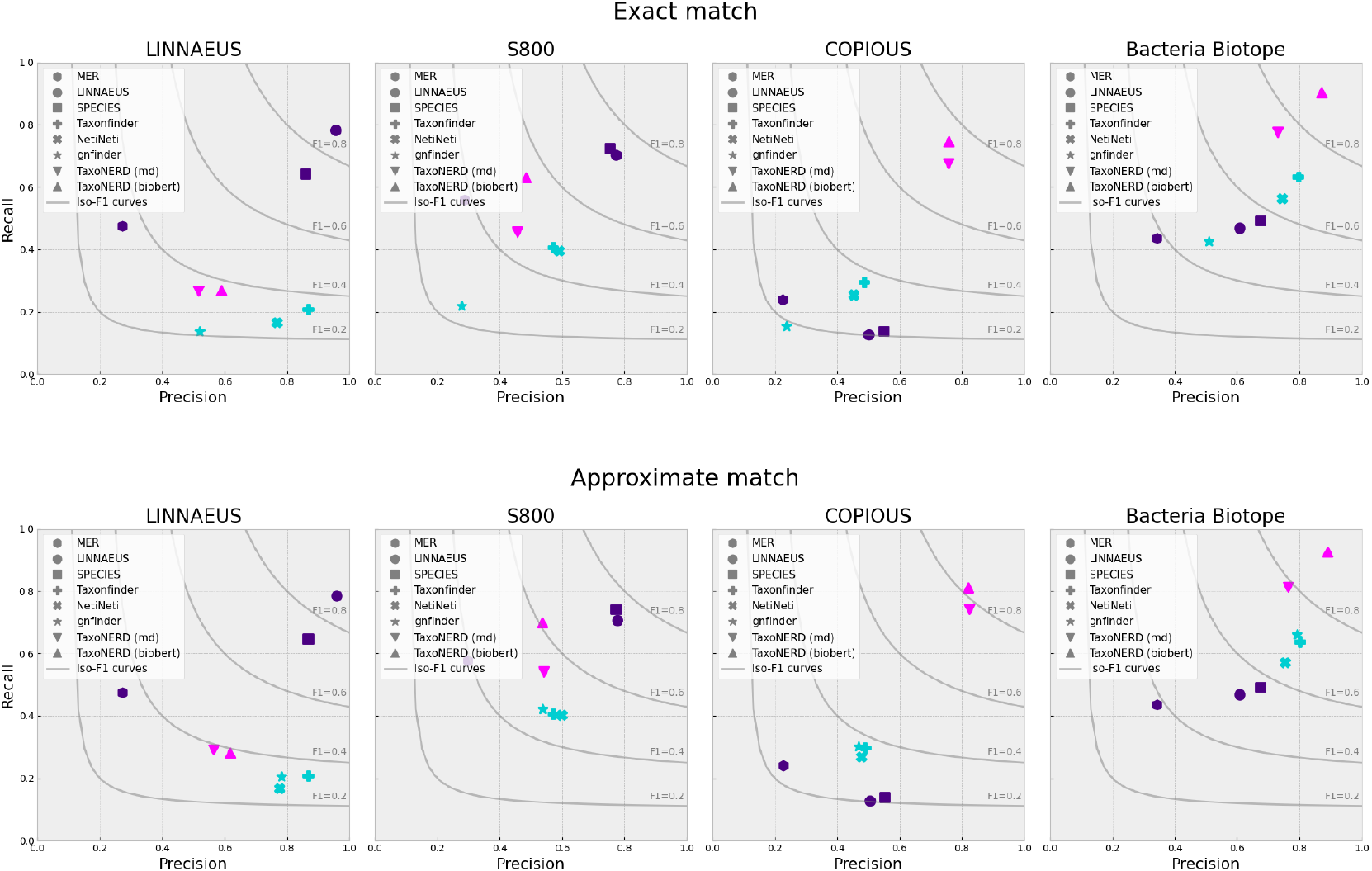
Precision and recall obtained on the four gold standard corpora using the exact match criterion (top) and the approximate match criterion (bottom). Grey lines represent iso-F_1_ curves. The different colors are used to distinguish between the different categories of tools : dictionary-based NER systems (violet), scientific name taggers (blue) and deep neural networks (pink).

**TABLE 3.** Precision, recall and F-score for the eight taxonomic NER systems evaluated on the four gold standard corpora, using exact match and approximate match as evaluation criteria.

| Corpus | Software | Exact match |  |  | Approximate match |  |  |
| --- | --- | --- | --- | --- | --- | --- | --- |
|  |  | PRE (%) | REC (%) | F<sub>1</sub> (%) | PRE (%) | REC (%) | F<sub>1</sub> (%) |
| LINNAEUS | MER | 27.35 | 47.40 | 34.69 | 27.38 | 47.40 | 34.71 |
|  | LINNAEUS | 95.71 | 78.14 | 86.04 | 96.03 | 78.39 | 86.32 |
|  | SPECIES | 86.04 | 64.22 | 73.55 | 86.73 | 64.73 | 74.13 |
|  | Taxonfinder | 86.83 | 20.84 | 33.61 | 86.83 | 20.84 | 33.61 |
|  | NetiNeti | 76.89 | 16.48 | 27.14 | 77.69 | 16.65 | 27.43 |
|  | gnfinder | 52.13 | 13.58 | 21.54 | 78.36 | 20.41 | 32.38 |
|  | TaxoNERD (md) | 51.75 | 26.56 | 35.10 | 56.57 | 29.08 | 38.42 |
|  | TaxoNERD (biobert) | 59.06 | 26.73 | 36.80 | 61.89 | 28.01 | 38.57 |
| S800 | MER | 28.82 | 55.80 | 38.01 | 29.86 | 57.63 | 39.34 |
|  | LINNAEUS | 77.41 | 70.14 | 73.60 | 77.84 | 70.53 | 74.01 |
|  | SPECIES | 75.31 | 72.36 | 73.80 | 77.20 | 74.19 | 75.66 |
|  | Taxonfinder | 57.04 | 40.68 | 47.49 | 57.04 | 40.68 | 47.49 |
|  | NetiNeti | 59.06 | 39.50 | 47.34 | 60.04 | 40.16 | 48.12 |
|  | gnfinder | 27.88 | 21.77 | 24.45 | 53.92 | 42.11 | 47.29 |
|  | TaxoNERD (md) | 45.74 | 45.50 | 45.62 | 54.26 | 54.05 | 54.15 |
|  | TaxoNERD (biobert) | 48.54 | 62.97 | 54.82 | 53.77 | 69.84 | 60.76 |
| COPIOUS | MER | 22.63 | 23.82 | 23.21 | 22.81 | 24.02 | 23.40 |
|  | LINNAEUS | 50.20 | 12.55 | 20.08 | 50.59 | 12.65 | 20.24 |
|  | SPECIES | 54.86 | 13.82 | 22.08 | 55.25 | 13.92 | 22.24 |
|  | Taxonfinder | 48.63 | 29.51 | 36.73 | 48.95 | 29.76 | 37.02 |
|  | NetiNeti | 45.36 | 25.39 | 32.56 | 47.81 | 26.82 | 34.36 |
|  | gnfinder | 23.85 | 15.29 | 18.64 | 46.94 | 30.10 | 36.68 |
|  | TaxoNERD (md) | 75.77 | 67.45 | 71.37 | 82.49 | 74.09 | 78.06 |
|  | TaxoNERD (biobert) | 75.85 | 74.51 | 75.17 | 82.14 | 81.00 | 81.57 |
| BB task | MER | 34.39 | 43.50 | 38.41 | 34.39 | 43.50 | 38.41 |
|  | LINNAEUS | 60.91 | 46.75 | 52.90 | 60.91 | 46.75 | 52.90 |
|  | SPECIES | 67.47 | 49.25 | 56.94 | 67.47 | 49.25 | 56.94 |
|  | Taxonfinder | 79.56 | 63.25 | 70.47 | 80.19 | 63.75 | 71.03 |
|  | NetiNeti | 74.50 | 56.25 | 64.10 | 75.50 | 57.00 | 64.96 |
|  | gnfinder | 51.05 | 42.50 | 46.38 | 79.28 | 66.00 | 72.03 |
|  | TaxoNERD (md) | 73.11 | 77.50 | 75.24 | 76.42 | 81.20 | 78.74 |
|  | TaxoNERD (biobert) | 87.20 | 90.25 | 88.70 | 89.13 | 92.48 | 90.77 |

Dictionary-based approaches (LINNAEUS and SPECIES) performed best on biomedically-oriented corpora (LINNAEUS and S800), with LINNAEUS achieving the highest F-score of all evaluated methods on the LINNAEUS corpus, and SPECIES being ranked first on the S800 corpus. This may be explained by the fact that the LINNAEUS and S800 corpora were first proposed as evaluation corpora for the LINNAEUS and SPECIES softwares respectively. It is likely that both methods were specifically tailored to perform well on their respective evaluation corpus. This could also explain the dramatic drop in performance observed for both methods on ecologically-oriented corpora (although the BB task corpus is composed of biomedical documents, it focuses on microorganisms ecology).

Scientific name taggers (Taxonfinder, NetiNeti, gnfinder) achieved a rather high precision on both the LINNAEUS and Bacteria Biotope task corpora (with Taxonfinder achieving the second higher precision rate on both corpora), at the exception of gnfinder whose precision using exact match is significantly lower than that of the other two methods. This can be attributed to the fact that gnfinder tends to significantly overestimate entity boundaries, including irrelevant punctuation marks and neighbouring words as part of the entities, which negatively impacts its performance using the exact match criterion. However, using approximate matching, the performance of gnfinder is similar to that of the other scientific name taggers. Their recall is consistently low on all corpora as these methods are designed to tag scientific names only, and all corpora also include annotations for common names. Despite using different approaches (dictionary, rules + machine learning, dictionary + machine learning), these methods perform quite similarly, with a slight advantage for Taxonfinder.

TaxoNERD’s deep neural models significantly outperformed all other approaches on the two ecological corpora (COPIOUS and the BB task corpus). Of our two DNN models, the Transformer-based model en_ner_eco_biobert consistently achieves the highest F-score on all corpora. TaxoNERD’s models show a tendency to slightly overestimate entity boundaries. A possible explanation is that COPIOUS annotations of taxon scientific names include the authorship information when available. As TaxoNERD’s models are partly trained on the COPIOUS corpus, they may have learned to recognise punctuation marks and other symbols following a taxon scientific name as being part of this name. Using the approximate match criterion that values boundaries overestimation, the F-score of our deep models increases by about 3-6%. The performance of TaxoNERD’s models on the COPIOUS corpora is of the same order of magnitude as that obtained by Nguyen et al. (2019) with their Bi-directional Long Short Term Memory (BiLSTM) model.

Finally, it is worth noting that our baseline dictionary-based approach (MER) performed poorly on all corpora, despite using a dictionary containing all the names in the NCBI Taxonomy. Nevertheless, its recall was significantly higher than that of LINNAEUS and SPECIES on the COPIOUS corpus.

## 4 DISCUSSION

The recognition of ecological concepts in text is a key technology for biodiversity information extraction and knowledge base population, and accurate NER tools are needed to make the most of the already large and continuously growing corpus of ecological texts. While researchers are mainly interested in species distribution, traits, habitats or interactions, a fundamental prerequisite to extract this information is the ability to detect mentions of taxa in text. A number of taxonomic NER tools are readily available to ecologists, including scientific name taggers and dictionary-based NER systems for recognising species names in biomedical documents.

Our experiments suggest that these existing tools are unsuitable to extract information from the ecological and evolutionary literature. Despite their relative effectiveness at detecting scientific names, as demonstrated on the BB task corpus which comprises a large proportion of such names, scientific name taggers are handicapped by their inability to detect common names. This is a barrier to ecological information extraction, as most references to taxonomic entities in the literature use their vernacular names, a taxon scientific name being often used only once or twice in the text for the sake of precision. The main alternative to scientific name taggers are dictionary-based taxonomic NER systems such as LINNAEUS and SPECIES. However, both methods rely on dictionaries that have been carefully tailored for biomedical use cases. As a result, these systems show high precision and recall on biomedical documents but their performance significantly drops on ecological corpora. When applied on ecological documents, both LINNAEUS and SPECIES miss many names, especially uninomials as these tools focus on species names. They are also prone to boundary estimation errors, missing authorship information and “sp.” or “spp.” abbreviations, as these forms of scientific names are not included in their dictionaries. Generally speaking, dictionary-based approaches lack robustness to previously unseen names, misspelling and other unexpected variants which are common in ecological papers. Their performance also depends heavily on the amount of effort put into creating a high-quality dictionary of names, as demonstrated by the poor performance of our baseline dictionary-based approach that uses a raw list of names from the NCBI Taxonomy.

None of the aforementioned taxonomic NER tools were able to recognise taxonomic entities in the COPIOUS corpus with satisfactory accuracy. COPIOUS being the biggest corpus of all four gold standards, it is also the one containing the largest diversity of taxon names, including scientific names (with or without authorship information), common names, abbreviations… which impedes the performance of dictionary-based NER systems and scientific name taggers. COPIOUS is also the only corpus composed exclusively of texts from the biodiversity literature, which causes biomedical NER tools like LINNAEUS and SPECIES to fail. TaxoNERD’s deep neural models on the contrary have been specifically trained to recognise taxon mentions in ecological documents and achieve high precision and recall on both COPIOUS and the BB task corpus. As anticipated, the Transfomer-based architecture achieves higher accuracy than spaCy’s standard architecture based on trigram CNNs. Although the difference in performance between the two models is quite spectacular on the BB task corpus (which could be attributed to the use of BioBERT, a language model pretrained on biomedical corpora, to obtain embeddings for words in the BB task corpus, composed of biomedical documents), this difference narrows on COPIOUS. This suggests that the en_ner_eco_md is also a good candidate for taxonomic NER in the ecological literature. The choice between the two models will depend on the compromise the user has to make between speed and accuracy, or on the availability of GPUs to speed up the inference process in Transformer-based models. Interestingly, the performance of TaxoNERD’s models on COPIOUS is consistent with the inter-annotator agreement reported by Nguyen et al. (2019) for this corpus. This suggests that our models are as good as human annotators at recognising taxon mentions in this corpus of documents.

Most of TaxoNERD’s errors on COPIOUS test set are due to TaxoNERD missing local (Filipino) vernacular names, and sometimes common English names (false negatives). Despite its propensity to overestimate entity boundaries, it also happens that TaxoNERD misses all or part of the authorship information (sometimes by a simple punctuation mark), which is punished twice by both criteria (one false positive, one false negative). On the BB task corpus, most errors are due to TaxoNERD missing alpha-numeric codes in strain names, e.g. “*B1157*” in “*L. lactis subsp. cremoris B1157*”, or to TaxoNERD’s tagging non-microorganism names, which are not annotated in the BB task corpus. Detecting strain mentions is recognised as a particularly difficult problem (Naderi et al., 2011) as they are prone to boundary estimation errors.

Although TaxoNERD is able to extract relevant information from ecological text with high precision and recall, performance drops on biomedical corpora. When looking at the predicted entities for the LINNAEUS and S800 corpora, we observe that TaxoNERD’s models tend to tag non-taxonomic scientific terms (“*Oligonucleoside methylphosphonates*”), allele and gene variant names, people names, and other capitalised expressions (“*Sequencing Kit*”, “*Staminal column*”, “*Immense tree*”) as taxonomic entities. Additionally, TaxoNERD failed to recognise srne terms considered as taxon names in the LINNAEUS and S800 corpora. This includes virus strain names and acronyms, such as “*H1N1*” or “”*H5 influenza virus*”, but also terms that do not directly convey species names such as “*patient*”, “*participants*” or “*people*”. As these terms are not relevant for ecological information extraction, this simply confirms that TaxoNERD is well suited for taxonomic NER in the ecological literature but should not be used (or at least carefully) for biomedical NER. It is also worth mentioning that the second major source of errors in the S800 corpus was the presence of many unannotated taxon names. Although TaxoNERD successfully detects these mentions, they are counted as false positives and result in a significant drop in precision.

DNN-based NER systems have achieved state-of-the-art results in a number of domains, and biomedical information extraction pipelines are now heavily relying on pretrained biomedical language models such as BioBERT (Lee et al., 2020), which are fine-tuned for downstream tasks, including named entity recognition and relation extraction. At the same time, we can see the number of initiatives to make DNN-based NLP solutions accessible to non experts multiplying. While researchers have access to state-of-the-art tools for biomedical information extraction (Perera et al., 2020), the ecological community has yet to get on the deep learning train and develop its own models and tools tailored to its specific use cases.

TaxoNERD is a very first step in this direction, which has not yet renounced its biomedical heritage, as it relies on pretrained biomedical models that are fine-tuned on an ecological corpus. Yet, it shows a significant gain in performance compared to existing tools for taxonomic NER in the biodiversity literature. Available as a command-line tool and a Python library, TaxoNERD can recognise all variants of scientific names and common names, as well as user-defined abbreviations thanks to scispaCy’s implementation of the simple abbreviation detection algorithm of Schwartz and Hearst (2002). With its two models, TaxoNERD provides different ways to balance speed and accuracy. TaxoNERD can also link taxon mentions to entities in a reference taxonomy using an approximate nearest neighbour search algorithm. Currently, TaxoNERD can link taxonomic entities to the NCBI Taxonomy (Federhen, 2002), GBIF Backbone Taxonomy (GBIF Secretariat, 2019) and TAXREF, the French national taxonomic register (Gargominy et al., 2019). Based on spaCy, TaxoNERD can be easily integrated into complex ecological information extraction pipelines while remaining very easy to use, bringing the predictive power of DNN-based NER systems to non-expert users.

TaxoNERD opens up many avenues for improving the performance of ecological NER systems. First, deep learning algorithms perform better with more data. To our knowledge, COPIOUS and the BB task corpus are the only gold standards designed specifically for ecological applications. Although this represents a significant amount of annotated documents, the performance of our models tends to peak. Using a larger training corpus, we could probably increase the accuracy of our models, or we could train more complex architectures with a greatest predictive power. However, as already mentioned, creating a GSC is a complex and costly process. A first alternative is data augmentation, which consists of expanding the training set by applying transformations to training instances without changing their labels (Dai and Adel, 2020). Another alternative is to use DNNs to learn a good language representation model from a large corpus of unannotated documents, and to use transfer learning to adapt the pretrained model to downstream tasks. In contrast to gold standards, large-scale unlabelled corpora are relatively easy to construct as they do not require any annotation effort. While there exists a number of pretrained language models for biomedical NLP, there exists none for ecological applications. Although we demonstrated with TaxoNERD that biomedical language models can be fine-tuned on ecological datasets with satisfactory performance, a domain-specific language representation pretrained on a large-scale ecological corpus would surely boost the performance of ecological information extraction tools.

Post-processing can also improve the quality and accuracy of predictions (Perera et al., 2020). For example, if a certain named entity is tagged once or several times in a document, and the same entity exist elsewhere in the text, untagged, then post-processing could make sure these missed NEs are also tagged with their predicted class, thus increasing recall. Another important subtask at this point is to resolve coreferences, i.e. mentions of taxa that appears as pronouns or noun phrases and which must be linked to the taxon names they refer to. Resolving coreferences is essential for a lot of higher-level information extraction tasks, including relation extraction, as much information concerning a taxon may be contained in sentences that do not explicitly use the taxon name. However, it is still considered one of the most difficult NLP tasks (Ng, 2017). State-of-the-art neural coreference resolution models have been made available in spaCy and require careful evaluation on ecological texts, but it seems likely that, as for many NLP tasks, domain-specific models will be needed to obtain better performance. Closely related to the problem of coreference resolution, entity normalisation is the task of disambiguating each textual mention to the correct entry in a given knowledge base. For instance, in the sentence “*Brown bears (Ursus arctos) flexibly change their feeding habits depending on the availability of dietary resources*”, the mentions “*Brown bears*”, “*Ursus arctos*” and “*their*” all refer to the unique entity with ID NCBI:txid9644 in the NCBI Taxonomy. Entity normalisation is a critical step in the process of turning unstructured textual information into machine-understandable facts.

In the longer term, we envision the creation a toolkit of state-of-the-art algorithms that would let ecologists and practitioners create their own pipelines to extract useful information on species distributions, traits or interactions from scientific and grey literature. This toolkit would include an ecological NER system with additional entity types (e.g. habitat, phenotype, etc.), coreference resolution and entity normalisation engines, a relation extractor to detect relationships between entities (e.g. interspecific interactions), and other NLP tools that will hopefully facilitate access to the considerable amount of knowledge held by the current (and future) body of published literature in ecology and evolution.

## acknowledgements

The research received funding from the French Agence Nationale de la Recherche (ANR) through the GlobNets (ANR-16-CE02-0009) project and through MIAI@Grenoble Alpes (ANR-19-P3IA-0003). We thank Lorraine Goeuriot and Martin Jeanmougin for their valuable comments and for proofreading the article.

## authors’ contributions

NLG and WT conceived the ideas and designed the study. NLG performed the literature review and the evaluation. NLG also designed TaxoNERD and trained the underlying models. NG produced the initial draft of the paper that was further revised and approved by WT.

## data availability

All the scripts used for the evaluation and the links to the four gold standard corpora are available on the GitHub repository accompanying this paper (https://github.com/nleguillarme/snr_tools_and_methods). TaxoNERD is available on PyPI under a MIT license; the sources, including the configuration files used to fine-tune the models, can be downloaded from the project’s GitHub repository (https://github.com/nleguillarme/taxonerd).

## Footnotes

1 https://github.com/nleguillarme/taxonerd

2 https://spacy.io/

3 https://allenai.github.io/scispacy/

4 https://github.com/spyysalo/s800

5 http://www.nactem.ac.uk/copious/

6 https://sites.google.com/view/bb-2019/

7 http://linnaeus.sourceforge.net/

8 https://species.jensenlab.org/

9 https://github.com/pleary/node-taxonfinder

10 https://github.com/dshorthouse/NetiNeti

11 https://github.com/gnames/gnfinder

12 http://ubio.org/index.php?pagename=namebank

13 https://github.com/nleguillarme/snr_tools_and_methods

